# Hippocampal network activity changes during early epileptogenesis predict subsequent epilepsy

**DOI:** 10.1101/2025.11.25.690377

**Authors:** Michael Strüber, Tristan Stöber, Vanessa Schütz, Lara Costard, Valentin Neubert, Sebastian Bauer, Julika Pitsch, Nico Melzer, Adam Strzelczyk, Jochen Triesch, Kai Siebenbrodt, Felix Rosenow, Ricardo Kienitz

**Affiliations:** Goethe University Frankfurt, Epilepsy Center Frankfurt Rhine-Main, Department of Neurology, University Hospital Frankfurt, Schleusenweg 2-16, 60528 Frankfurt am Main, Germany; LOEWE Center for Personalized Translational Epilepsy Research (CePTER), Goethe University Frankfurt, Frankfurt am Main, Germany; Institute for Neural Computation, Ruhr University Bochum, Bochum, Germany; Eli Lilly Cork Ltd., Island House, Eastgate Business Park, Little Island, Co. Cork, Ireland; Oscar Langendorff Institute of Physiology, University of Rostock, Rostock, Germany; Department of Epileptology, University Hospital Bonn, Bonn, Germany; Department of Neurology, Medical Faculty and University Hospital Düsseldorf, Heinrich Heine University of Düsseldorf, Düsseldorf 40225, Germany; Frankfurt Institute for Advanced Studies, Ruth-Moufang-Straße 1, 60438 Frankfurt am Main, Germany

## Abstract

The circuit mechanisms underlying focal epileptogenesis are, despite of decades of epilepsy research, still incompletely understood. In this study, we aimed to characterize the changes in hippocampal network activity induced by a potentially epileptogenic insult. In rats, long-lasting electrical perforant pathway stimulation leads in a high percentage of animals to the development of temporal lobe epilepsy. However, a subset of animals remains resilient against the stimulation. We monitored alterations of neuronal activity by chronically recording the local field potential (LFP) from the hippocampal dentate gyrus before, during and after the potentially epileptogenic insult. Intriguingly, *epilepsy* animals identified by subsequent spontaneous epileptic seizures were characterized by a transient increase in the aperiodic exponent suggesting a shift towards a reduced local excitation-to-inhibition (E/I) ratio during the first days after the perforant path stimulation. Furthermore, these animals developed a strong impairment of theta oscillation prevalence and regularity during early epileptogenesis. In contrast, resilient *non-epilepsy* animals without spontaneous seizures neither showed this modulation in E/I ratio nor a corruption of hippocampal theta activity. In fact, the increase in the aperiodic exponent on the first day after completion of the electrical stimulation paradigm could predict epileptogenesis with very high fidelity (AUC 0.92) and correlated significantly with later seizure rate. This finding opens the opportunity to dissect mechanisms of epileptogenesis and to test the effectiveness of anti-epileptogenesis treatment in very early disease stages by allowing identification of individuals at high risk. Furthermore, it might offer a potential explanation for the frequently observed failure of anti-epileptogenesis drugs boosting GABAergic inhibition.

## Introduction

The hallmarks of epilepsy as one of the most prevalent chronic neurological diseases are dysfunctional cortical circuits and epileptic seizures^1^, generated by sudden changes in brain network activity. This excessive and highly synchronized neuronal activity causes severe paroxysmal symptoms such as loss of consciousness, falls, involuntary body movements or vegetative dysregulation with apnea or asystole in extreme cases. On the level of brain networks, the exact mechanisms of seizure generation but also the alterations underlying the propensity of neuronal networks to generate seizure activity are only partially understood. Inhibitory interneurons are prominent regulators of neuronal activity and play an important role for brain function in health and disease^2^. Their inhibitory action can counterbalance excessive excitation in neuronal networks but at the same time they exert a synchronizing effect on their target cell population^3,4^. Thus, it is not surprising that data on their pro- or anti-epileptic role are ambiguous and sometimes contradictory^5^.

The changes occurring during the development of acquired focal epilepsy, the so-called *epileptogenesis* period, are complex and are thought to involve neuronal, glial, immune- and blood-brain-barrier related mechanisms^1,6–8^. However, essential for the strong propensity of brain tissue to generate seizures are alterations in neuronal networks rendering them more excitable^9^. One critical determinant of neuronal network excitability is the ratio of excitatory and inhibitory synaptic activity distributed in the circuit, the so-called excitation/inhibition-(E/I-) ratio^10^. The time-dependent changes of E/I ratio in neuronal networks during their epileptogenic conversion to hyperexcitable circuits have not been addressed.

In the last years, novel methods have been developed to estimate the amount of inhibition in neuronal networks by analysis of local field potential (LFP) recordings^11,12^. In this study, we monitored changes in hippocampal network excitability over the course of epileptogenesis in a toxin-free model of electrically-induced temporal lobe epilepsy in rats.

## Materials and Methods

### Animals and electrode positioning

All animal experiments were conducted in accordance with national and institutional guidelines (EU directive 2010/63/EU; approved by the Regierungspräsidium Gießen and the Regierungspräsidium Darmstadt). Altogether 18 adult Sprague-Dawley male rats (weight 300-450 g) were housed individually to avoid damage of implants by littermates. Animals had free access to food and drinking water, a 12h/12h light/dark cycle was maintained. Six of the 18 included animals have already delivered data to a previous study^13^.

The surgical procedure has been previously described in detail^13,14^. Briefly, stimulation electrodes (0.125 mm in diameter) were implanted bilaterally in the angular bundle of the perforant path (coordinates: ±4.5 mm lateral to bregma, 0.5 mm rostral to lambda, variable depth). Furthermore, a recording electrode (0.25 mm in diameter) was implanted unilaterally in the left hippocampus (coordinates relative to bregma: 2 mm lateral, 3 mm caudal, variable depth). Over three consecutive days, electrical stimulation was applied over 30 min on the first two days and over 8 hours on the third stimulation day. A more detailed description of the surgery and epileptogenesis induction protocol can be obtained from the **Supplementary Material**, the electrical stimulation pattern is shown in **Fig. 1A**.

**Figure 1.**
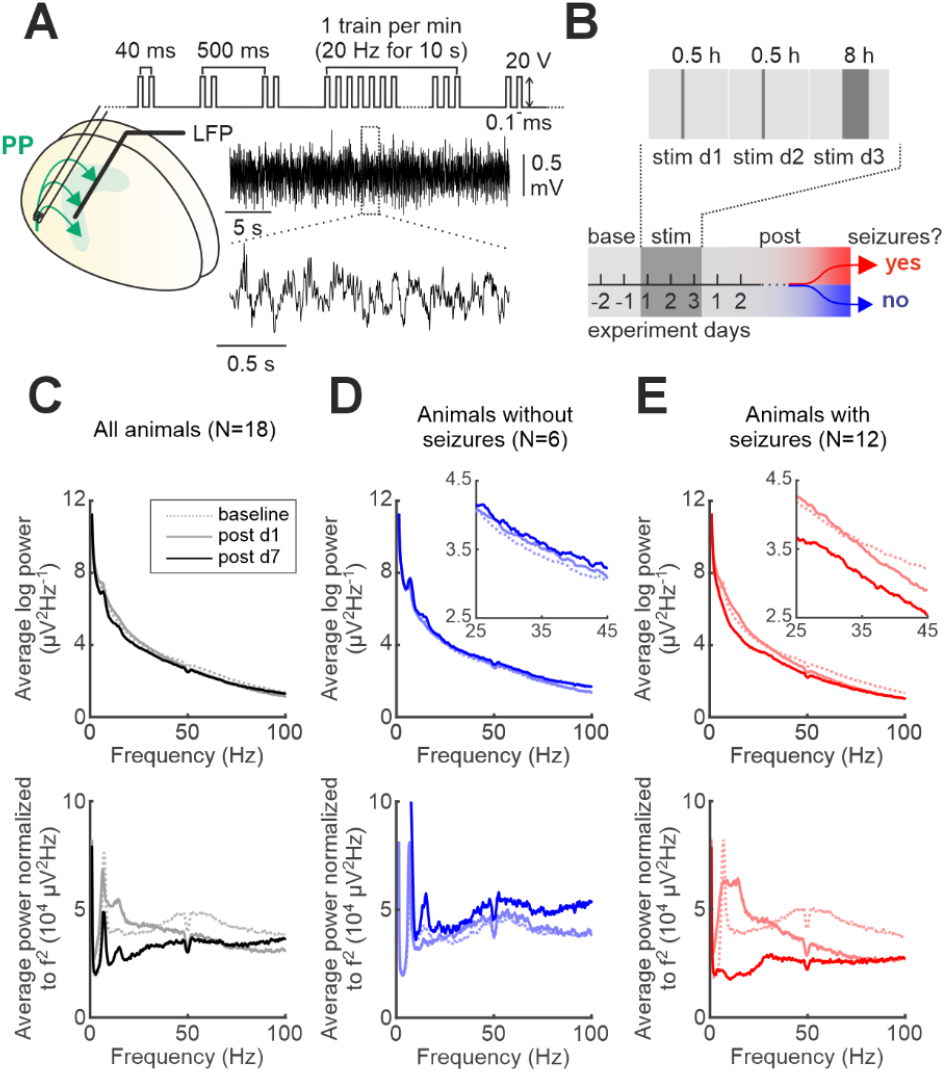
Electrically induced early epileptogenesis. **A** Experimental setup with bilateral perforant path (PP) stimulation and continuous local field potential (LFP) recordings from hippocampal dentate gyrus. Top, stimulation protocol; middle, representative recorded LFP; bottom, magnified episode with prevalent theta oscillations. **B** Time line of the experiment is schematically shown. On 3 stimulation days, the stimulation protocol shown in A is applied over 30 min on stim days 1 and 2 and over 8 h on stim d3. After completion of stimulation, most animals develop spontaneous epileptic seizures after ~21 days (epilepsy) while a minority remained free from spontaneous seizures (non-epilepsy). **C-E** Power density analysis of the LFP recorded during baseline (light color, dashed line) and on 1st (light color, continuous line) and 7th day (dark color) after stimulation averaged over all animals (**C**, black), over all non-epilepsy animals (**D**, blue) and over all epilepsy animals (**E**, red). Top, power plotted after logarithmic transformation; bottom, power is frequency-normalized. Insets, focus on the frequency range 25-45 Hz. Note, specifically on post d1 the slope of logarithmic power spectrum is increased in epilepsy animals.

### Analysis of electrophysiological data

All data analysis was performed in MATLAB (The Mathworks, version R2022a). To prevent the results from being biased by electrical artifacts, EEG snippets devoid of artifacts with a duration of 2 min were manually selected for further analysis. To characterize long-term changes during the course of the experiment independent from daytime, for every analysis day EEG snippets were isolated at 3, 4 and 5 am and the respective results averaged. For the analysis of the time course around the individual stimulation sessions with a higher temporal resolution, again three individual 2 min EEG snippets were manually selected at pre-defined latencies from the end of the respective stimulation (i.e. 30min, 1h, 2h, 4h, 8h and 18h). As a baseline for every stimulation session, the EEG snippets of the respective stimulation day collected in the morning hours (3-5 am) were used. Stimulation usually commenced at 11 am-1 pm of the respective day.

To remove 50 Hz power net artifacts, EEG signals were in a first step bandpass-filtered using a power spectrum interpolation filter (*dft* filter of the fieldtrip toolbox^15^). Power spectra were obtained using the *mtspectrumc* function of the Chronux toolbox (http://chronux.org). Analysis of the aperiodic activity and determination of the aperiodic exponent was performed using the *fooof*-algorithms of the fieldtrip toolbox^15^ with analysis over the frequency range 1-95 Hz. However, similar results have been obtained by performing the analysis over the frequency range 25-45 Hz.

### Statistics

Statistical significance was tested using two-tailed versions of the indicated tests. Distributions were tested for normality using Lilliefors’ test (MATLAB’s *lillietest*). Whenever the normality hypothesis was rejected, a Wilcoxon rank sum test was used. Otherwise an unpaired two-sample T test was employed.

Values in the text are given as mean±standard error of the mean (SEM), if not stated otherwise. To analyze correlations, Spearman rank analysis was used.

To investigate the influence of outcome (*epilepsy* vs *non-epilepsy*) on the prevalence of significant theta oscillation activity, repeated measures ANOVA was used (MATLAB’s *fitrm* and *ranova*). The p-value given in the text describes the interaction between outcome and time (**Fig. 3A**).

To estimate fidelity of epileptogenesis prediction on the basis of the aperiodic exponent change upon perforant path stimulation in 18 animals, a linear discriminant classification model was used in a leave-one-out cross validation approach. In detail, we generated 18 training sets each consisting of 17 animals with a different animal left out in each classification round. In every round, a linear discriminant classification model using MATLAB’s *fitcdiscr* function was trained on the training data and tested with the data of the animal left out in the respective round. Receiver operating characteristics (ROC) analysis (**Fig. 2F**) was performed on the 18 prediction scores resulting from the cross validation procedure using MATLAB’s *rocmetrics* function. No oversampling was performed due to only mild class imbalance (12 versus 6 observations).

**Figure 2.**
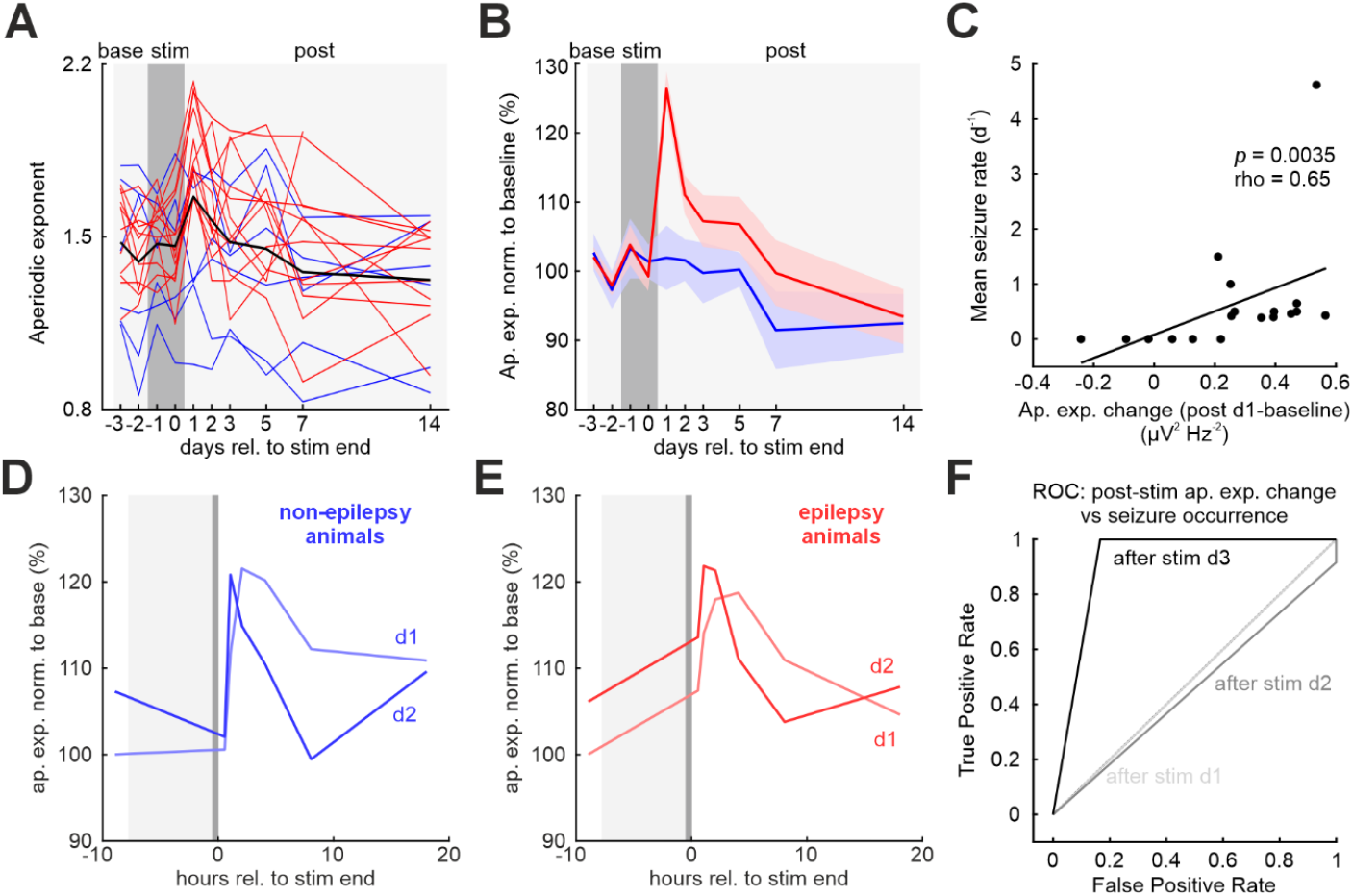
Transient reduction of E/I ratio during initial epileptogenesis. **A** Time course of the aperiodic exponent as a proxy for the local E/I ratio before, during and after electrical stimulation over three days (dark grey area). Lines represent individual non-epilepsy (blue, 6 animals) and epilepsy animals (red, 12 animals). Thick black line, average over all 18 animals. **B** Mean time course of the aperiodic exponent during the experiment in non-epilepsy (blue) and epilepsy (red) animals. Lines, mean; shaded areas, SEM. Epilepsy but not non-epilepsy animals show a transient increase of the aperiodic exponent lasting several days after completed stimulation (p=0.014, repeated measures ANOVA). **C** Post-stimulation increase of the aperiodic exponent correlates significantly with the chronic seizure rate. p, rho, Spearman rank correlation analysis. **D**,**E** After stimulation on days 1 and 2, the aperiodic exponent on average increases for a period of a few hours in both, non-epilepsy (blue, D) and epilepsy (red, E) animals. **F** Receiver operating characteristics (ROC) analysis illustrating the predictive value of the aperiodic exponent change after stimulations for the classification of animals as non-epilepsy or epilepsy (after stim d1, AUC 0.5; after stim d2, AUC 0.46; after stim d3, AUC 0.92).

**Figure 3.**
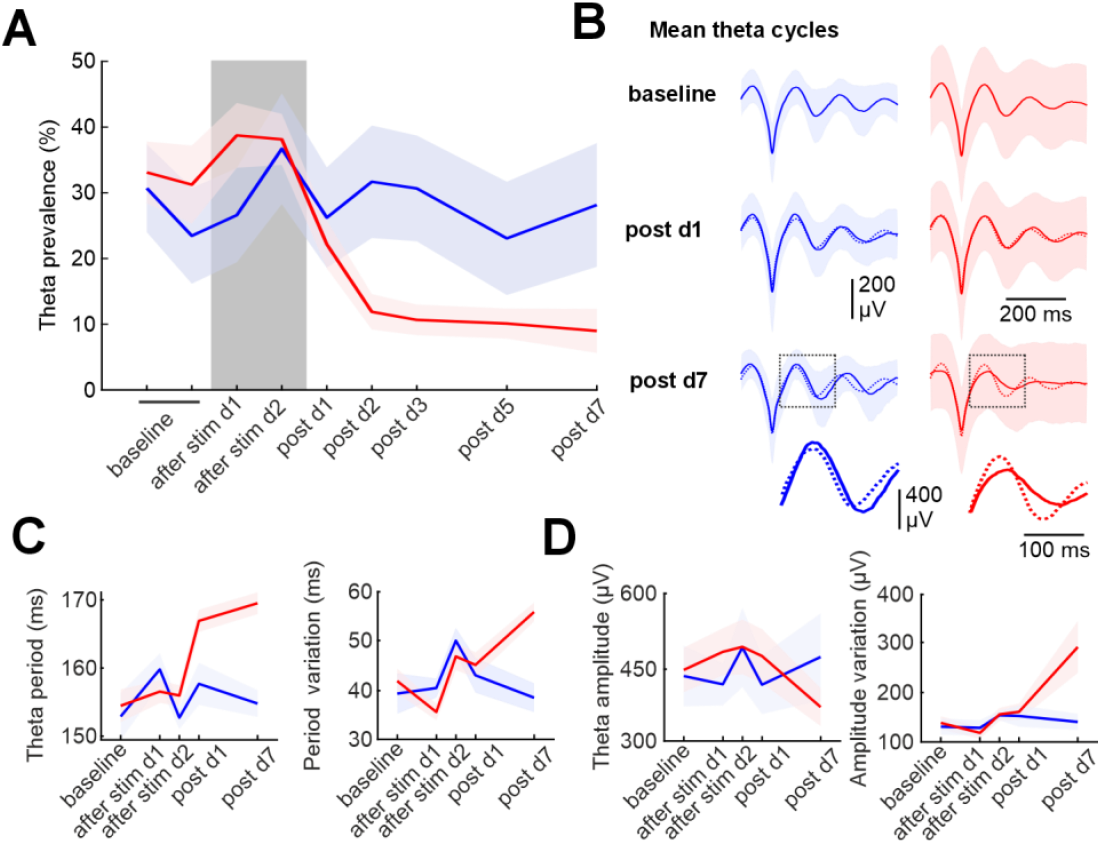
Reduced theta prevalence and regularity during early epileptogenesis. **A** Percentage of analyzed epochs classified as theta-dominated for non-epilepsy (blue) and epilepsy (red) animals at different time points of the experiment. **B** Mean theta oscillation waveform aligned to the theta trough and averaged over all non-epilepsy (left, blue) and epilepsy (right, red) animals at baseline, 1^st^ and 7^th^ day after stimulation. Continuous lines, average theta waveform; dashed lines, baseline waveform. Bottom, magnification of average theta waveforms indicated by dashed boxes. **C**,**D** Average theta cycle properties at different time points: period duration (C, left); variation of period duration of consecutive cycles (C, right); amplitude (D, left); variation of amplitude of consecutive cycles (D, right). Lines, mean; shaded areas, standard deviation (A,B) or SEM (C,D).

## Results

In response to the long-lasting electrical hippocampal stimulation over three consecutive days modelling electrically induced epileptogenesis (**Fig. 1B; Materials and Methods**), 12 of 18 included animals developed spontaneous seizures after a mean latency of 21.4±3.1 days (*epilepsy* rats) and 6 showed no spontaneous electrographical seizures during the observation period (*non-epilepsy* rats). Six of 18 animals showed acute symptomatic seizures defined as seizures occurring in the first 7 days after completion of stimulation^16^ (33% prevalence of acute symptomatic seizures in *epilepsy* and *non-epilepsy* rats, respectively; see **Supplementary Fig. S1**). We characterized changes in hippocampal network activity induced by the perforant path stimulation by power spectrum density analysis (**Fig. 1C-E**). Recent methodological advances suggested the inference of the ratio of excitatory and inhibitory synaptic activity in neuronal circuits from the 1/f^*χ*^ relationship of aperiodic LFP signal power, with *χ* being coined the *aperiodic exponent*^12^. While higher ratios of aperiodic excitatory-to-inhibitory (E/I) activity are reflected by flatter power spectra particularly for the sub-gamma frequency range (~<40-50 Hz), lower E/I ratios are associated with more steeply declining power spectra^11,12^. Surprisingly, *epilepsy* animals but not *non-epilepsy* animals showed more steeply declining power spectra in the frequency range of 25-45 Hz on the first day after stimulation compared to both baseline and post 7d recordings (**Fig. 1 D,E insets**).

To further investigate this finding, we determined for every animal the aperiodic exponent at multiple time points before, during and after stimulation. On average, the aperiodic exponent seemed to increase strongly after the third, 8h lasting stimulation but was reduced to baseline levels already on day 7 after stimulation (**Fig. 2A**). When analyzing *epilepsy* and *non-epilepsy* rats separately, a prominently elevated aperiodic exponent upon stimulation was only found in those rats developing spontaneous seizures during the following weeks (**Fig. 2B**, *p*=0.014, repeated measurements ANOVA). In support of this finding, we found a significant positive correlation between chronic seizure rate and the absolute increase in the aperiodic exponent on the first day after completion of the stimulation (**Fig. 2C**). Next, we checked whether the shorter stimulations on days 1 and 2 could induce shifts in the E/I ratio, too. We monitored the aperiodic exponent at a higher temporal resolution around the stimulation time points and found a substantial but brief increase following the 30 min stimulation in both, *epilepsy* and *non-epilepsy* animals (**Fig. 2D,E**; see also **Supplementary Fig. S2**). Thus, highly intense 30 min-lasting perforant pathway stimulation shifts the E/I ratio towards lower values for hours. However, long lasting stimulation over 8 h seems to induce a much more stable state of inhibitory dominance in animals developing chronic epilepsy. Finally, we performed a receiver operating characteristics analysis of a linear discriminant classification model trained on the data of all 18 animals and found that the change in the aperiodic exponent after the third stimulation day but not after the first or the second stimulation day could predict subsequent development of epilepsy with high fidelity (AUC = 0.92; **Fig. 2F**).

To further characterize hippocampal network function, we studied the properties of theta oscillations, the most prominent oscillatory pattern in the hippocampus strongly associated with animal locomotion^17^. To avoid any behavior-dependent bias, we selected LFP epochs with significant oscillatory activity in the theta frequency range (5-9 Hz) for further analysis (see **Supplementary Materials and Methods**). The average fraction of LFP epochs showing significant theta activity was not different in *non-epilepsy* and *epilepsy* animals during baseline conditions and after stimulation days 1 and 2. However, after the final stimulation on day 3, *epilepsy* animals showed hippocampal theta activity during significantly less recording time than *non-epilepsy* rats (*p*=0.021, repeated measures ANOVA; **Fig. 3A**). Next, we focussed only on LFP epochs containing significant theta oscillatory activity (cf **Supplementary Fig. S3**). To detect changes in the theta oscillation characteristics, we obtained mean theta cycle waveforms aligned to the oscillatory trough averaged over all *non-epilepsy* and *epilepsy* animals (**Fig. 3B**). During the course of early epileptogenesis, average theta cycles seemed to be horizontally stretched and vertically compressed in *epilepsy* rats. In order to dissect the underlying alterations in oscillation structure, we performed a cycle-by-cycle analysis of theta activity at different time points (**Fig. 3C,D**). Theta cycle period showed a sudden increase upon completion of the stimulation pattern already on *post d1* in *epilepsy* but not *non-epilepsy* animals. However, theta cycle amplitude was not altered by the stimulation protocol. Next, we probed stability of theta oscillations and studied the cycle-to-cycle variability in both period duration and amplitude of theta activity. For both, cycle amplitude and period duration a marked increase in the cycle-to-cycle variability became evident for *epilepsy* but not *non-epilepsy* animals on day 7 after stimulus completion (**Fig. 3C,D**). Thus, early epileptogenesis after electrical perforant path stimulation is characterized by a substantially reduced regularity of theta oscillations.

In the hippocampal dentate gyrus, theta activity temporally structures excitatory and inhibitory synaptic activity with afferent excitation from the entorhinal cortex arriving at granule cells shortly before the theta trough with a mean theta phase angle of −39° ^18^. Indeed, when pooling all animals during baseline conditions and determining the aperiodic exponent at different theta phases, the aperiodic exponent showed a clear theta phase-dependent modulation with a minimum at −30° (**Fig. 4A**), indicating maximal glutamatergic synaptic activity shortly before the theta trough as shown by Pernía-Andrade and Jonas^18^. To test epileptogenesis related alterations in the temporal structure of synaptic excitation and inhibition, we quantified the theta phase dependent modulation of the aperiodic exponent at *post d1* and *post d7* in *epilepsy* and *non-epilepsy* animals (**Fig. 4B**). While in the *non-epilepsy* animals the aperiodic exponent displayed at all analyzed time points a clear minimum related to the theta trough, this phase modulation was inverted in *epilepsy* animals on *post d1* with a maximum shortly after the theta trough. On *post d7*, theta phase modulation of the aperiodic exponent returned to baseline levels with a minimum at the time point of theta troughs. Thus, the reduction in E/I ratio in early epileptogenesis seems to be most prominent during the theta trough when afferent perforant path input has been shown to excite the dentate gyrus circuit.

**Figure 4.**
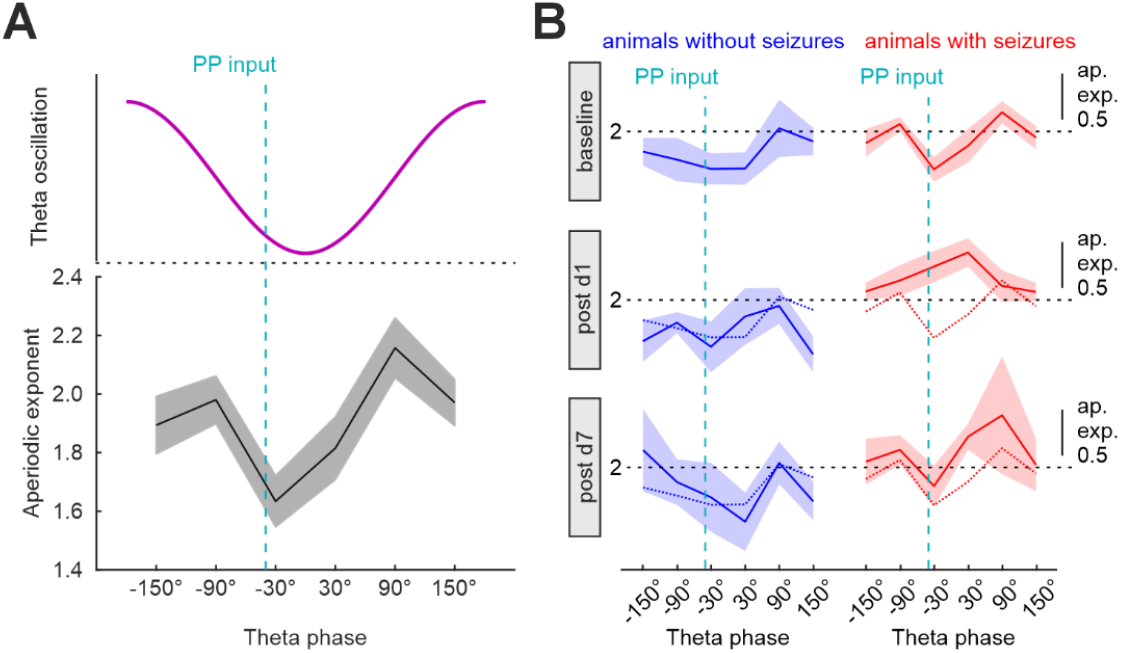
Epileptogenesis-associated elevation of inhibitory dominance is theta phase-dependent. **A** Theta phase-dependent modulation of the aperiodic exponent with a minimum at −30° corresponding well to the phase of maximal perforant path-mediated excitation (cyan broken line). Black line, average over all 18 animals during baseline; shaded area, SEM. **B** Evolution of the theta phase dependent aperiodic exponent after perforant path stimulation in non-epilepsy (blue, left) and epilepsy (red, right) animals. Continuous lines, average over all animals at the respective experimental day; broken lines, baseline; shaded areas, SEM.

## Discussion

The neuronal circuit mechanisms underlying the progressively developing propensity of epileptic networks to generate seizures are to date far from being fully understood. Synaptic inhibition provided by GABAergic interneurons is known to play an ambiguous role in the generation of seizures with both, seizure-suppressive and seizure-igniting effects^5^. Here, we show in an electrically induced temporal lobe epilepsy rat model the transient increase in the relative abundance of inhibition immediately after application of an epileptogenic incident specifically in animals found to subsequently develop epilepsy (**Fig. 2**). This transient post-incidental dominance of circuit inhibition could actually predict the development of epilepsy with very high fidelity and might therefore be classified as a potential electrophysiological marker for early epileptogenesis.

Whether the observed change in E/I ratio is indeed due to elevated inhibitory synaptic activity, a reduced excitatory neurotransmission or both is not clear. Furthermore, the cellular processes underlying the alteration in E/I ratio remain to be determined in future studies. The longer time scale of the E/I ratio shift towards inhibition lasting over several days suggests that potential mechanisms lie beyond mere extracellular ion concentration shifts or different time courses of relief from refractory ion channels. In fact, changes in the E/I ratio in response to variations in the input excitation to neuronal networks have been described in various *in vitro* and *in vivo* systems and are subsumed under the concept of *homeostatic plasticity*^19,20^. Thus, we hypothesize, that extensive electrical perforant path stimulation could induce homeostatic plasticity processes such as synaptic scaling or adaptation of cellular excitability finally leading to a longer lasting shift in the local E/I ratio^21^. The tight temporal association of the E/I reduction with the time point of perforant path-mediated excitation of the dentate gyrus network (**Fig. 4B**) suggests that the potentiation of feedforward or feedback inhibitory circuits are responsible for the observed effects. Although at this stage, discussing possible specific plasticity mechanisms remains mere speculation, the sole enhancement of the excitability of GABAergic interneurons providing feedforward and feedback inhibition to granule cells could fully explain the observed epileptogenesis-related alteration of the local E/I ratio. Indeed, for hippocampal Parvalbumin-positive (PV^+^) interneurons different plasticity mechanisms leading to elevated excitability have been described: Intense synaptic excitation of PV+ interneurons in the dentate gyrus can enhance intrinsic excitability for prolonged periods of time by shifting the resting membrane potential to more depolarized levels^22^. Furthermore, Donato and colleagues described different excitability states for PV^+^ interneurons with a highly excitable state corresponding to a high level of Parvalbumin- and GAD67-expression being induced by strong excitation of the interneuron network^23^.

Whether the observed transient relative dominance of inhibitory tone acts protective for the hippocampal circuits, as it could be *a priori* assumed, or alternatively contributes to epileptogenesis itself, needs to be further clarified. One argument in favour of a causal role of early inhibitory dominance for epilepsy development lies in the exclusive observation of the transient inhibition increase in *epilepsy* animals. A presumable protective mechanism could be expected to be more strongly expressed in those animals undergoing the potentially epileptogenic incident but remaining resilient against it. In agreement with this line of argumentation, maladaptive homeostatic plasticity has already been discussed as potential driving factors for ictogenesis and epileptogenesis^24–26^. In fact, recently, temporal lobe epilepsy patients were shown to express higher aperiodic exponents in seizure-generating networks than healthy counterparts^27^. Mechanistically, a delayed, GABA-triggered and microglia-mediated elimination of GABAergic synapses on principal cells has been recently identified in the hippocampus during kainic acid induced epileptogenesis and hypothesized to be one mechanism of established hyperexcitability mediated by an increased E/I ratio at later stages^28^.

Post-insult transient changes in the E/I ratio may not only play a causative role in the generation and progression of epilepsy, but could in principle also underlie functional deficits in epileptic brain circuits. The impairment of brain network function in chronic epilepsy patients is one of the dominant symptoms beside seizures^29^. In our data set, theta oscillations become less regular with progressing epileptogenesis (**Fig. 3C,D**). As theta oscillations particularly in rodents are well known to serve as a reference signal for hippocampal information processing, the identified destruction of theta regularity might plausibly contribute to hippocampal dysfunction. Changes in the E/I ratio with elevated inhibition have been shown to, in general, affect brain theta oscillations in a similar way (reduced theta frequency and power)^30^ and to, more specifically, impair hippocampal theta-dependent neuronal coding^31^.

### Technical limitations

One of the most evident points of criticism lies in the estimation of the ratio of inhibitory and excitatory synaptic activity by analyzing the aperiodic component of LFP power spectra. Since the distribution of this method by Voytek and colleagues^11^, many neuroscientists have applied this easy-to-use methodology to address diverse highly relevant scientific questions. Although valid criticism has been brought forward including the recently detected intrusion of cardiac signals in aperiodic activity recorded from the brain^32^, many example situations of evident shifts in the E/I ratio are captured by according changes in the aperiodic exponent of the power spectra of respective neuronal signals^11,12,30,33–35^. Thus, while the analysis of aperiodic activity remains an indirect proxy for the actual ratio of excitatory and inhibitory synaptic activity in brain circuits, many lines of evidence indicate its usefulness in gross E/I ratio estimation. Indeed, our reproduction of the theta-phase dependent rise in excitation shortly before the theta trough stands in very good agreement with published data^18^ and further confirms the validity of our E/I ratio-estimation by aperiodic network activity analysis. Nevertheless, the exact cellular mechanisms underlying the changes in hippocampal network activity during early epileptogenesis need to be investigated in much more detail in future studies. Moreover, future research needs to assess, whether a transient phase of inhibitory dominance is a general phenomenon of early epileptogenesis comprising both animal epilepsy models of different etiology and human epilepsy development.

## Conclusions

In an electrically induced temporal lobe epilepsy model in rats, the process of epileptogenesis hidden to date can be electrophysiologically revealed as early as on the first day after the application of the epileptogenic stimulus by determining the time course of the aperiodic exponent of the hippocampal LFP. This finding has several potential implications for our understanding of epileptogenesis. Firstly, network activity in epileptogenic circuits is drastically altered already one day after the incident^36^. Secondly, in analogy to the generation of epileptic seizures, synaptic inhibition might not only act protective during the process of epileptogenesis but may even drive the development of epileptic circuits at least during the initial days. Finally, these results open the opportunity to dissect the mechanisms of protection from epileptogenesis by identifying future epileptic animals already immediately after stimulus application with a very high fidelity – questions inevitably related to the long-held hope for an anti-epileptogenesis treatment in high risk patients suffering from potentially epileptogenic conditions such as intracranial trauma or new-onset status epilepticus.

## Author Contributions

MS, TS, RK, JT, AS and FR contributed to the conception and design of the study. MS, TS, VS, LC, VN, JP, KS and SB collected and analyzed data. MS, TS, NM and RK wrote the manuscript.

## References

1. Devinsky, O., Vezzani, A., O’Brien, T.J., Jette, N., Scheffer, I.E., de Curtis, M., and Perucca, P. (2018). Epilepsy. Nat. Rev. Dis. Primer 4, 1–24. 10.1038/nrdp.2018.24.

2. Pelkey, K.A., Chittajallu, R., Craig, M.T., Tricoire, L., Wester, J.C., and McBain, C.J. (2017). Hippocampal GABAergic Inhibitory Interneurons. Physiol. Rev. 97, 1619–1747. 10.1152/physrev.00007.2017.

3. Sessolo, M., Marcon, I., Bovetti, S., Losi, G., Cammarota, M., Ratto, G.M., Fellin, T., and Carmignoto, G. (2015). Parvalbumin-Positive Inhibitory Interneurons Oppose Propagation But Favor Generation of Focal Epileptiform Activity. J. Neurosci. Off. J. Soc. Neurosci. 35, 9544–9557. 10.1523/JNEUROSCI.5117-14.2015.

4. de Curtis, M., and Avoli, M. (2016). GABAergic networks jump-start focal seizures. Epilepsia 57, 679–687. 10.1111/epi.13370.

5. Wenzel, M., Huberfeld, G., Grayden, D.B., de Curtis, M., and Trevelyan, A.J. (2023). A debate on the neuronal origin of focal seizures. Epilepsia 64 Suppl 3, S37–S48. 10.1111/epi.17650.

6. Vezzani, A., Balosso, S., and Ravizza, T. (2019). Neuroinflammatory pathways as treatment targets and biomarkers in epilepsy. Nat. Rev. Neurol. 15, 459–472. 10.1038/s41582-019-0217-x.

7. Vezzani, A., Ravizza, T., Bedner, P., Aronica, E., Steinhäuser, C., and Boison, D. (2022). Astrocytes in the initiation and progression of epilepsy. Nat. Rev. Neurol. 18, 707–722. 10.1038/s41582-022-00727-5.

8. Batulin, D., Lagzi, F., Vezzani, A., Jedlicka, P., and Triesch, J. (2022). A mathematical model of neuroimmune interactions in epileptogenesis for discovering treatment strategies. iScience 25, 104343. 10.1016/j.isci.2022.104343.

9. Liou, J.-Y., Smith, E.H., Bateman, L.M., Bruce, S.L., McKhann, G.M., Goodman, R.R., Emerson, R.G., Schevon, C.A., and Abbott, L.F. (2020). A model for focal seizure onset, propagation, evolution, and progression. eLife 9, e50927. 10.7554/eLife.50927.

10. Shao, L.-R., Habela, C.W., and Stafstrom, C.E. (2019). Pediatric Epilepsy Mechanisms: Expanding the Paradigm of Excitation/Inhibition Imbalance. Children 6, 23. 10.3390/children6020023.

11. Gao, R., Peterson, E.J., and Voytek, B. (2017). Inferring synaptic excitation/inhibition balance from field potentials. NeuroImage 158, 70–78. 10.1016/j.neuroimage.2017.06.078.

12. Donoghue, T., Haller, M., Peterson, E.J., Varma, P., Sebastian, P., Gao, R., Noto, T., Lara, A.H., Wallis, J.D., Knight, R.T., et al. (2020). Parameterizing neural power spectra into periodic and aperiodic components. Nat. Neurosci. 23, 1655–1665. 10.1038/s41593-020-00744-x.

13. Costard, L.S., Neubert, V., Venø, M.T., Su, J., Kjems, J., Connolly, N.M.C., Prehn, J.H.M., Schratt, G., Henshall, D.C., Rosenow, F., et al. (2019). Electrical stimulation of the ventral hippocampal commissure delays experimental epilepsy and is associated with altered microRNA expression. Brain Stimulat. 12, 1390–1401. 10.1016/j.brs.2019.06.009.

14. Norwood, B.A., Bumanglag, A.V., Osculati, F., Sbarbati, A., Marzola, P., Nicolato, E., Fabene, P.F., and Sloviter, R.S. (2010). Classic hippocampal sclerosis and hippocampal-onset epilepsy produced by a single “cryptic” episode of focal hippocampal excitation in awake rats. J. Comp. Neurol. 518, 3381–3407. 10.1002/cne.22406.

15. Oostenveld, R., Fries, P., Maris, E., and Schoffelen, J.-M. (2011). FieldTrip: Open source software for advanced analysis of MEG, EEG, and invasive electrophysiological data. Comput. Intell. Neurosci. 2011, 156869. 10.1155/2011/156869.

16. Pitkänen, A., and Bolkvadze, T. (2012). Head Trauma and Epilepsy. In Jasper’s Basic Mechanisms of the Epilepsies, J. L. Noebels, M. Avoli, M. A. Rogawski, R. W. Olsen, and A. V. Delgado-Escueta, eds. (National Center for Biotechnology Information (US)).

17. Sławińska, U., and Kasicki, S. (1998). The frequency of rat’s hippocampal theta rhythm is related to the speed of locomotion. Brain Res. 796, 327–331. 10.1016/S0006-8993(98)00390-4.

18. Pernía-Andrade, A.J., and Jonas, P. (2014). Theta-gamma-modulated synaptic currents in hippocampal granule cells in vivo define a mechanism for network oscillations. Neuron 81, 140– 152. 10.1016/j.neuron.2013.09.046.

19. Turrigiano, G.G. (2017). The dialectic of Hebb and homeostasis. Philos. Trans. R. Soc. B Biol. Sci. 372, 20160258. 10.1098/rstb.2016.0258.

20. Lee, H.-K., and Kirkwood, A. (2019). Mechanisms of Homeostatic Synaptic Plasticity in vivo. Front. Cell. Neurosci. 13. 10.3389/fncel.2019.00520.

21. Buhl, E.H., Otis, T.S., and Mody, I. (1996). Zinc-induced collapse of augmented inhibition by GABA in a temporal lobe epilepsy model. Science 271, 369–373. 10.1126/science.271.5247.369.

22. Ross, S.T., and Soltesz, I. (2001). Long-term plasticity in interneurons of the dentate gyrus. Proc. Natl. Acad. Sci. U. S. A. 98, 8874–8879. 10.1073/pnas.141042398.

23. Donato, F., Rompani, S.B., and Caroni, P. (2013). Parvalbumin-expressing basket-cell network plasticity induced by experience regulates adult learning. Nature 504, 272–276. 10.1038/nature12866.

24. Lignani, G., Baldelli, P., and Marra, V. (2020). Homeostatic Plasticity in Epilepsy. Front. Cell. Neurosci. 14. 10.3389/fncel.2020.00197.

25. Issa, N.P., Nunn, K.C., Wu, S., Haider, H.A., and Tao, J.X. (2023). Putative roles for homeostatic plasticity in epileptogenesis. Epilepsia 64, 539–552. 10.1111/epi.17500.

26. Savin, C., Triesch, J., and Meyer-Hermann, M. (2009). Epileptogenesis due to glia-mediated synaptic scaling. J. R. Soc. Interface 6, 655–668. 10.1098/rsif.2008.0387.

27. Duma, G.M., Cuozzo, S., Wilson, L., Danieli, A., Bonanni, P., and Pellegrino, G. (2024). Excitation/Inhibition balance relates to cognitive function and gene expression in temporal lobe epilepsy: a high density EEG assessment with aperiodic exponent. Brain Commun. 6, fcae231. 10.1093/braincomms/fcae231.

28. Chen, Z.-P., Zhao, X., Wang, S., Cai, R., Liu, Q., Ye, H., Wang, M.-J., Peng, S.-Y., Xue, W.-X., Zhang, Y.-X., et al. (2025). GABA-dependent microglial elimination of inhibitory synapses underlies neuronal hyperexcitability in epilepsy. Nat. Neurosci. 28, 1404–1417. 10.1038/s41593-025-01979-2.

29. Vinti, V., Dell’Isola, G.B., Tascini, G., Mencaroni, E., Cara, G.D., Striano, P., and Verrotti, A. (2021). Temporal Lobe Epilepsy and Psychiatric Comorbidity. Front. Neurol. 12.

30. Salvatore, S.V., Lambert, P.M., Benz, A., Rensing, N.R., Wong, M., Zorumski, C.F., and Mennerick, S. (2024). Periodic and aperiodic changes to cortical EEG in response to pharmacological manipulation. J. Neurophysiol. 131, 529–540. 10.1152/jn.00445.2023.

31. Valero, M., Navas-Olive, A., de la Prida, L.M., and Buzsáki, G. (2022). Inhibitory conductance controls place field dynamics in the hippocampus. Cell Rep. 40, 111232. 10.1016/j.celrep.2022.111232.

32. Schmidt, F., Danböck, S.K., Trinka, E., Klein, D.P., Demarchi, G., and Weisz, N. (2024). Age-related changes in “cortical” 1/f dynamics are linked to cardiac activity. eLife 13. 10.7554/eLife.100605.1.

33. Brake, N., Duc, F., Rokos, A., Arseneau, F., Shahiri, S., Khadra, A., and Plourde, G. (2024). A neurophysiological basis for aperiodic EEG and the background spectral trend. Nat. Commun. 15, 1514. 10.1038/s41467-024-45922-8.

34. Diehl, G.W., and Redish, A.D. (2024). Measuring excitation-inhibition balance through spectral components of local field potentials. BioRxiv Prepr. Serv. Biol., 2024.01.24.577086. 10.1101/2024.01.24.577086.

35. Pochinok, I., Stöber, T.M., Triesch, J., Chini, M., and Hanganu-Opatz, I.L. (2024). A developmental increase of inhibition promotes the emergence of hippocampal ripples. Nat. Commun. 15, 738. 10.1038/s41467-024-44983-z.

36. Bumanglag, A.V., and Sloviter, R.S. (2018). No latency to dentate granule cell epileptogenesis in experimental temporal lobe epilepsy with hippocampal sclerosis. Epilepsia 59, 2019–2034. 10.1111/epi.14580.

