## Supplementary Information for "Hippocampal network activity changes during early epileptogenesis predict subsequent epilepsy"

### Supplementary Figures

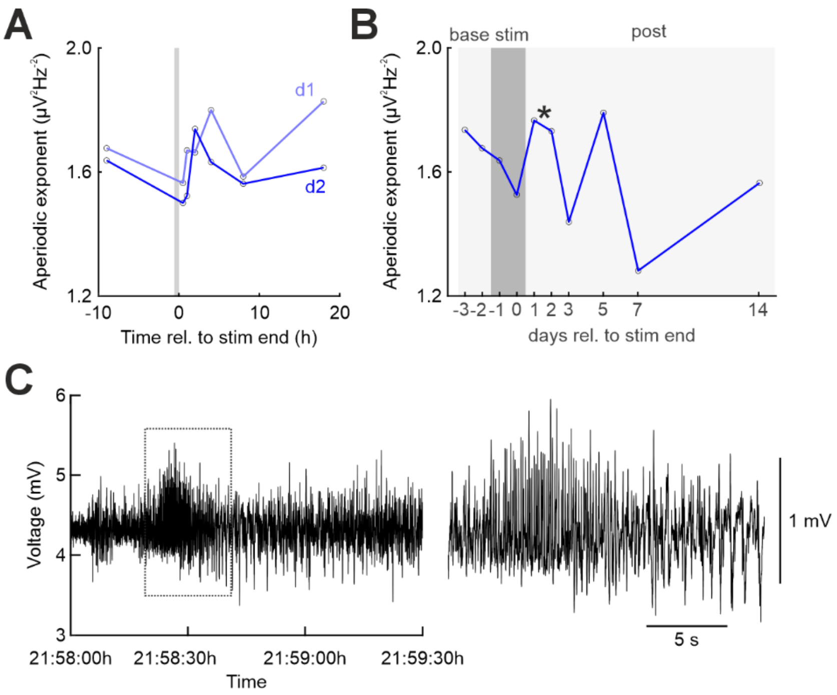


**Supplementary Figure S1. Non-epilepsy animal #139 shows strong response to electrical perforant path stimulation but no transient increase in the aperiodic exponent. A** Time course of the aperiodic exponent before and immediately after the stimulations on stim d1 (light blue) and on stim d2 (dark blue). Stimulation period is indicated by light grey rectangle. Note the clear, brief post-stimulation increase in the aperiodic exponent after completion of stimulation. **B** Time course of the aperiodic exponent before, during and after the 3 days-stimulation session. Note the absence of a transient, days-long increase in the aperiodic exponent after completion of the complete stimulation session. Stimulation days are indicated by dark grey rectangle. Asterisk, time point of acute symptomatic seizure depicted in C. **C** LFP recording during a representative acute symptomatic seizure on the first day after completion of the stimulation session. LFP trace in the dotted box is magnified on the right.

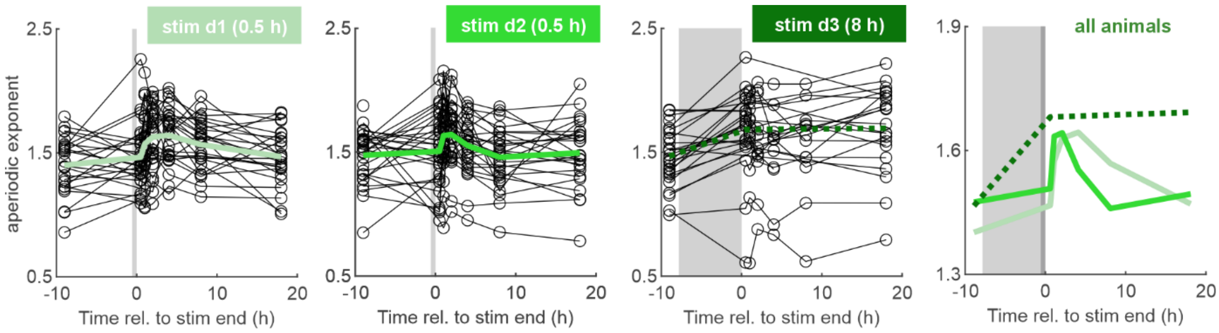


**Supplementary Figure S2. Brief period of elevated aperiodic exponent immediately after stimulus application.** Time course of the aperiodic exponent in the hours before and immediately after electrical perforant path stimulation on stimulation days 1, 2 and 3 (from left to right). Grey boxes, stimulation periods; circles connected by lines, individual animals; thick green lines, mean aperiodic exponent time course averaged over all included animals. Note, only a reduced number of animals could be analyzed on stim d3 due to lasting seizures patterns and afterdischarges in the respective analysis time window. Rightmost, average aperiodic exponent time course over all animals after stimulation on days 1 (light green), 2 (green continuous line) and 3 (dark green, dashed line).

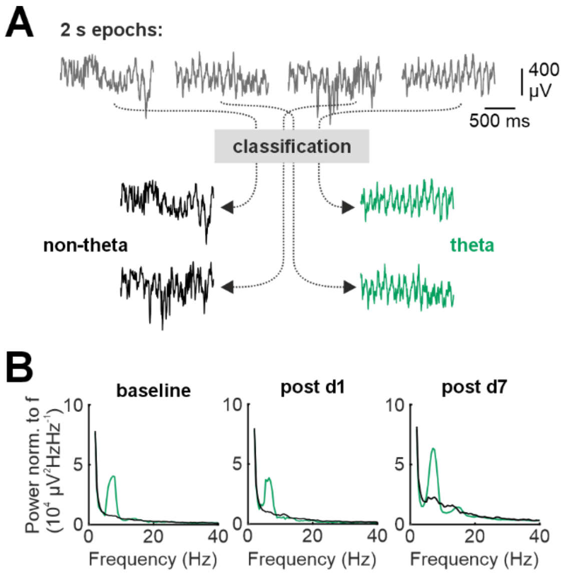


**Supplementary Figure S3. Isolation of significant theta epochs. A** Epochs of 2 s long local field potential (LFP) epochs were classified as either non-theta (black) or theta (green) dominated LFP epoch (see Materials and Methods). **B** Representative example average power spectra of one animal (#143) obtained from recordings before (baseline, left), 1 days (post d1, middle) and 7 days (post d7, right) after completed stimulation based on the non-theta (black) and the theta (green) epochs.

### Supplementary Materials and Methods

*Surgery, epileptogenesis induction and seizure detection*

After initial implantation, depths of recording and stimulation electrodes were adjusted during repetitive electrical stimulation to maximize the amplitude of the dentate gyrus field population spike in response to perforant path stimulation. After the adjustments, implants were fixed to the skull by dental cement including additional fixation and ground screws inserted in the bone. To obtain continuous EEG recordings, the implanted recording electrode and ground cables were subcutaneously connected to an EEG wireless transmitter (OpenSource Instruments [OSI], Waltham, MA, USA), implanted in a subcutaneous pouch on the right flank. After surgery, animals were allowed to recover for seven days before the beginning of perforant path stimulation.

At the beginning of each of the three stimulation days, animals were briefly anesthetized with isoflurane and the electrical stimulator was bilaterally connected to the stimulation electrodes. The parameters of the stimulation paradigm are illustrated in **Fig. 1A**. Rats were preconditioned on stimulation days 1 and 2 by delivering 30 min of repetitive current pulses as a preparation for the final epileptogenesis induction over 8 hours on stimulation day 3[1]. During stimulation, LFP was recorded from the dentate gyrus in order to guarantee efficient stimulation. Immediately after stimulation completion, animals were briefly anesthetized with isoflurane to avoid and interrupt potential self-sustained status epilepticus and put back to their home cages where chronic intracranial EEG recordings recommenced (bandpass filter 0.3-160 Hz, sampling rate 512 Hz) for several weeks (observation time after stimulation 54.5±9.0 days, mean±SEM) to detect the occurrence, the latency to and the rate of spontaneous seizures. The electrophysiological data were analyzed in total and screened for seizure patterns. The occurrence of seizures was detected electrographically and confirmed clinically by inspection of continuously recorded videos by at least two reviewers with high expertise in EEG reading. Seizures occurring within the first seven days of recording were defined as *acute symptomatic seizures*.

*Theta oscillation analysis*

To identify significant theta-episodes, a two-step procedure was applied. First, 2min EEG snippets were bandpass-filtered in the range 4-9 Hz (2nd order Butterworth filter) and split in 2s-long epochs. For all epochs the respective envelope signal and its mean ($\mu_{env}$) were determined (MATLAB's *envelope* function). Epochs were classified as significant theta epochs, if:

Eq. (1) $\mu_{env} > 0.5 \cdot \sigma(V_{4\text{-}9Hz})$

with $\sigma$, standard deviation; $V_{4\text{-}9Hz}$, 4-9 Hz-bandpass-filtered LFP

All other 2s-epochs were classified as *non-theta*.

In a second step, for all unfiltered 2s-LFP epochs the relative fraction of the 4-9 Hz power among the total power in the 2-40 Hz range ($\theta$rel) was calculated. Then, for every animal, a specific relative theta power-threshold ($th_{\theta rel}$) was defined as the 95-percentile of the distribution of relative theta power $\theta$rel of all *non-theta* 2s-epochs. Only significant theta-epochs with a relative theta-power > $th_{\theta rel}$ were included in subsequent analyses.

Individual theta cycles within the theta epochs were isolated by identifiying zero-crossings and determining the respective peaks and troughs. Average theta cycle waveforms were obtained by generating trough-aligned averages of the isolated theta cycles over all animals in the respective groups (**Fig. 3B**). Period durations of individual cycles were determined by the time between subsequent troughs, theta cycle amplitudes were defined as the difference between theta peaks and the mean of the adjacent troughs (**Fig. 3C,D**). Cycle-to-cycle variation of period duration and amplitude were defined as the absolute difference between period and amplitude of a given cycle and the overnext cycle.

### Supplementary References

1. Norwood, B. A. *et al.* Classic hippocampal sclerosis and hippocampal-onset epilepsy produced by a single 'cryptic' episode of focal hippocampal excitation in awake rats. *J. Comp. Neurol.* **518**, 3381–3407 (2010).
